# Shades of grey: Coat-colour dependent effect of urbanization on the bacterial microbiome of a wild mammal

**DOI:** 10.1101/2021.02.16.431540

**Authors:** Mason R. Stothart, Amy E.M. Newman

**Affiliations:** Department of Ecosystem and Public Health, Faculty of Veterinary Medicine, University of Calgary, Calgary T2N 4Z6, Canada; Department of Integrative Biology, College of Biological Sciences, University of Guelph, Guelph N1G 2W1, Canada

**Keywords:** colour polymorphism, dispersal limitation, eastern grey squirrel, gene x environment interactions, microbial ecology, null modelling, plasticity, 16S rRNA gene

## Abstract

**Background:** Host-associated microbiota can be fundamental to the ecology of their host and may even help wildlife species colonize novel niches or cope with rapid environmental change. Urbanization is a globally replicated form of severe environmental change which we can leverage to better understand wildlife microbiomes. Does the colonization of separate cities result in parallel changes in the intestinal microbiome of wildlife, and if so, does within-city habitat heterogeneity matter? Using 16S rRNA gene amplicon sequencing, we quantified the effect of urbanization on the microbiome of eastern grey squirrels (*Sciurus carolinensis*). Eastern grey squirrels are ubiquitous in both rural and urban environments throughout their native range, across which they display an apparent coat colour polymorphism (agouti, black, intermediate).

**Results:** Grey squirrel microbiomes differed between rural and city environments; however, comparable variation was explained by habitat heterogeneity within cities. Our analyses suggest that operational taxonomic unit (OTU) community structure was more strongly influenced by local environmental conditions (rural and city forests versus human built habitats) than urbanization of the broader landscape (city versus rural). Many of the bacterial genera identified as characterizing the microbiomes of built-environment squirrels are though to specialize on host-derived products and have been linked in previous research to low fibre diets. However, despite an effect of urbanization at fine spatial scales, phylogenetic patterns in the microbiome were coat colour phenotype dependent. City and built environment agouti squirrels displayed greater phylogenetic beta-dispersion than those in rural or forest environments, and null modelling results indicated that the phylogenetic structure of urban agouti squirrels did not differ greatly from stochastic phylogenetic expectations.

**Conclusions:** Habitat heterogeneity at fine spatial scales affects host-associated microbiomes, however, we found little evidence that this pattern was the result of similar selective pressures acting on the microbiome within environments. Further, this result, those of phylogeny-independent analyses, and patterns of beta-dispersion lead us to suggest that microbiota dispersal and ecological drift are integral to shaping the inter-environmental differences we observed. These patterns were partly mediated by squirrel coat colour phenotype, and therefore putatively, host physiology. Given a well-known urban cline in squirrel coat colour melanism, grey squirrels provide an ideal free-living system with which to study how host genetics mediate environment x microbiome interactions.

## Background

Recognition that host-associated microbial communities (microbiomes) affect host health [1, 2], phenotypes [3, 4], and fitness [5, 6] has sparked interest in understanding the causes, consequences, and eco-evolutionary relevance of microbiome variation in nature [7–10]. The microbiome’s greater mutability (relative to the host genome) undergirds theorization that the microbiome plays an important evolutionary role in vertebrates [10–12]. Namely, it is hypothesized that the microbiome may help to buffer the adverse affects of novel environmental change by extending a host’s phenotypic range. In doing so, the microbiome might facilitate adaptive stop-gap solutions during host colonization to a new ecological niche. Central to this hypothesis is the prediction that among host populations, there exists genetic variation in traits that shape microbiome plasticity [11]. Many free-living study systems are intractable for empirically testing these predictions, partly because it is difficult to define what constitutes a truly ‘novel’ environmental change.

Urban ecosystems have achieved recognition in the field of evolutionary biology as opportunistic, mass-replicated, free-living ecological and evolutionary experiments [13, 14]— experiments which (we argue) are useful for probing the host-microbiome relationship. Cities are among the fastest growing terrestrial ecosystems in the world [15] and expose wildlife to combinations of stressors and stimuli unlike anything observed in nature [16]. The novelty of urban environmental selective pressures is such that, globally, rates of phenotypic change among plants and animals show a distinct urban signature [17]. However, for many of these phenotypic changes, an underlying host genetic basis has not be strongly demonstrated [18]. The rapidity of phenotypic change of plants and animals in response to urbanization, and the nascence of cities on an evolutionary timescale, make urban environments a useful testing-ground for hypotheses that the microbiome can facilitate wildlife or—from a holobiont perspective—drive adaptation to a new ecological niche [11,19,20].

Despite widespread evidence for an effect of urbanization on the microbiomes of wild plants and animals, reported patterns are inconsistent across host species [21–29]. Inconsistencies of urban effects are likely attributable to; 1) city-specific variation (most studies to-date have focused on populations residing within a single city); 2) differences in the operational definition of urbanization; 3) differences in the how host species interact with the urban environment. For example, among vertebrates, consumption of human food resources is hypothesized to be the primary driver of microbiome variation [24,26,30,31]. Dietary variation is likely an important contributor to urban microbiome variation for some wildlife, but not all species rely on human food subsidies to the same extent [32]. Alongside diet, patterns of intra- and inter-specific microbiota dispersal between hosts may help shape microbiome differences between environments [33–36]. Likewise, the physiological responses of hosts to environmental stimuli may be more important than the details of the stimuli itself [29, 37]. The capacity of the microbiome to facilitate wildlife expansion to a new ecological niche may therefore strongly depend on the outcome of gene x environment x microbiome interactions [38].

Building on past research, we adopt a multi-city perspective in studying the effect of urbanization on the faecal microbiome of eastern grey squirrels (*Sciurus carolinensis*)—a species with a prominent coat colour polymorphism which is both linked to urbanization and pleiotropically connected to gene x environment interactions among squirrels [39, 40]. Specifically, this colour polymorphism is the result of an incompletely dominant mutation in the gene which encodes the melanocortin-1 receptor of the proopiomelanortin system [41]; agouti: homozygous for the wild type allele, black: homozygous for the mutant allele, intermediate: heterozygous for the wild-type and mutant alleles (Figure 1). Along replicate rural-urban gradients, the melanic phenotype has been reported to increase in frequency relative to the agouti wild-type [42, 43]. Fur and feather colour polymorphisms are frequently pleiotropically linked to differences in vertebrate physiology and behaviour, such as immunity, stress physiology, aggression, and even grooming [44]. Similar coat colour linked differences among squirrels could explain the relative success of melanic phenotypes on the urban landscape and may mediate how the squirrel microbiome responds to urban environmental challenges.

**Figure 1:**
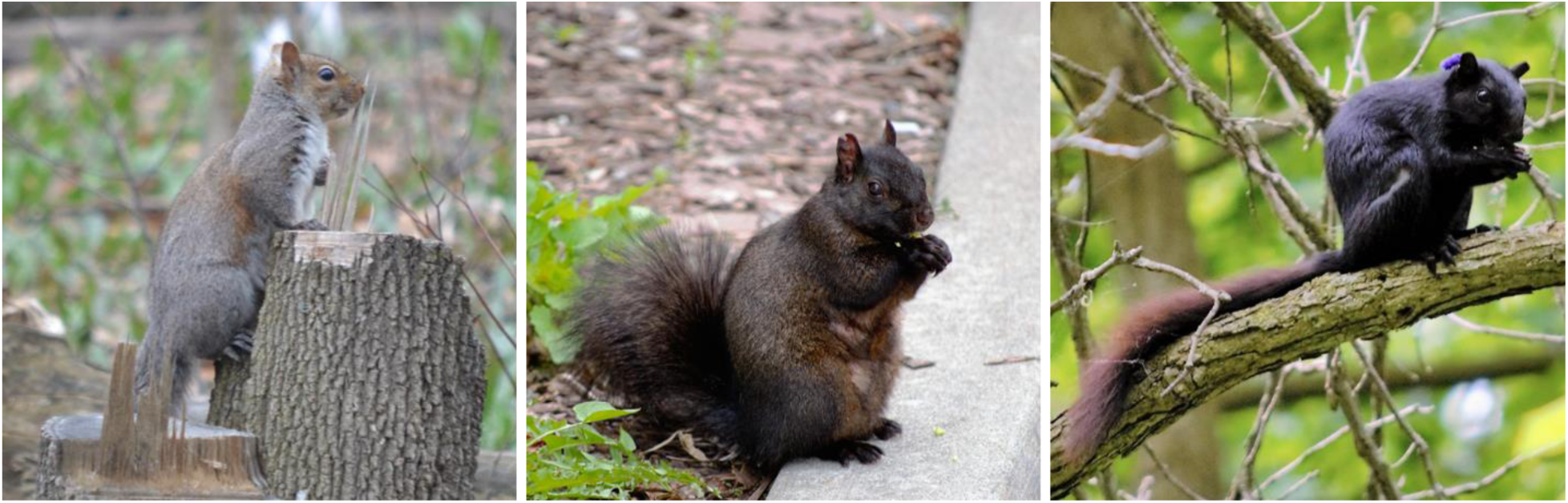
Representative photos of the (left) agouti (zero copies of the melanism causing MC1R-Δ24 mutant allele), (centre) intermediate (one copy of the MC1R-Δ24 mutant allele), and (right) black (two copies of the MC1R-Δ24 mutant allele) eastern grey squirrel (*Sciurus carolinensis*) coat colour phenotypes [41].

Using a 16S rRNA gene amplicon sequencing approach, we characterized the faecal bacterial microbiome of 195 individual adult eastern grey squirrels spanning three rural forests and three cities across southern Ontario (Canada), including three distinct urban land-classes within the city of Guelph. As most urban wildlife microbiome research relies on measurements made within a single city, we tested the prediction that the independent colonization of separate cities by eastern grey squirrels has resulted in paralleled changes in the bacterial microbiome. To inferentially parse what (non-mutually exclusive) ecological processes (e.g., selection, dispersal, drift) might underlay inter-environmental microbiome variation, we used a phylogenetic null modelling approach [45–47]. We predicted that homogenizing selection should act on the microbiome within environments, but that evidence for disparate selective pressures acting on the microbiome would be observed between environments. To evaluate competing hypotheses that local (forest versus human built environment) [21, 23] versus landscape (city versus rural) [25] level environmental variation was more important in shaping the microbiome, we contrasted variation in the microbiome explained by between-versus within-city habitat classifications (university campuses, suburban parks, urban forests, rural forests). Finally, to test for an effect of intra-specific physiological variation, we tested for interactions between coat-colour phenotype and the bacterial microbiome’s response to urbanization.

## Results

### α-diversity

We observed 1256 operational taxonomic units (OTUs) across 195 squirrel faecal samples, with an average of 379 OTUs ± 4 *SE* per squirrel. The eastern grey squirrel microbiome was comprised primarily of *Lachnospiraceae* (37% ± 1% *SE*), *Ruminococcaceae* (19% ± 0.6% *SE*), *Muribaculaceae* (10% ± 0.5% *SE*), *Prevotellaceae* (8% ± 0.5% *SE*), *Lactobacillaceae* (7% ± 0.8% *SE*), and *Bacteroidaceae* (5% ± 0.5% *SE*). No difference in α-diversity was observed between sexes (*β* = 2.11 ± 8.12 *SE*, *t* = 0.26, *p* = 0.80). Agouti squirrels from city sites trended towards having fewer OTUs than agouti squirrels from rural sites, albeit this effect was marginally non-significant (*β* = 59.88 ± 31.17 *SE*, *t* = −1.921, *p* = 0.06). However, we observed a colour phenotype x environment interaction, whereby α-diversity decreased with increases in MC1R mutation number (agouti: 0, intermediate: 1, black: 2) at rural sites (*β* = −29.39 ± 12.26 *SE*, *t* = −2.40, *p* = 0.02) but increased with melanism mutation number at city sites (*β* = 37.27 ± 13.86 *SE*, *t* = 2.69, *p* < 0.01; Figure 2).

**Figure 2:**
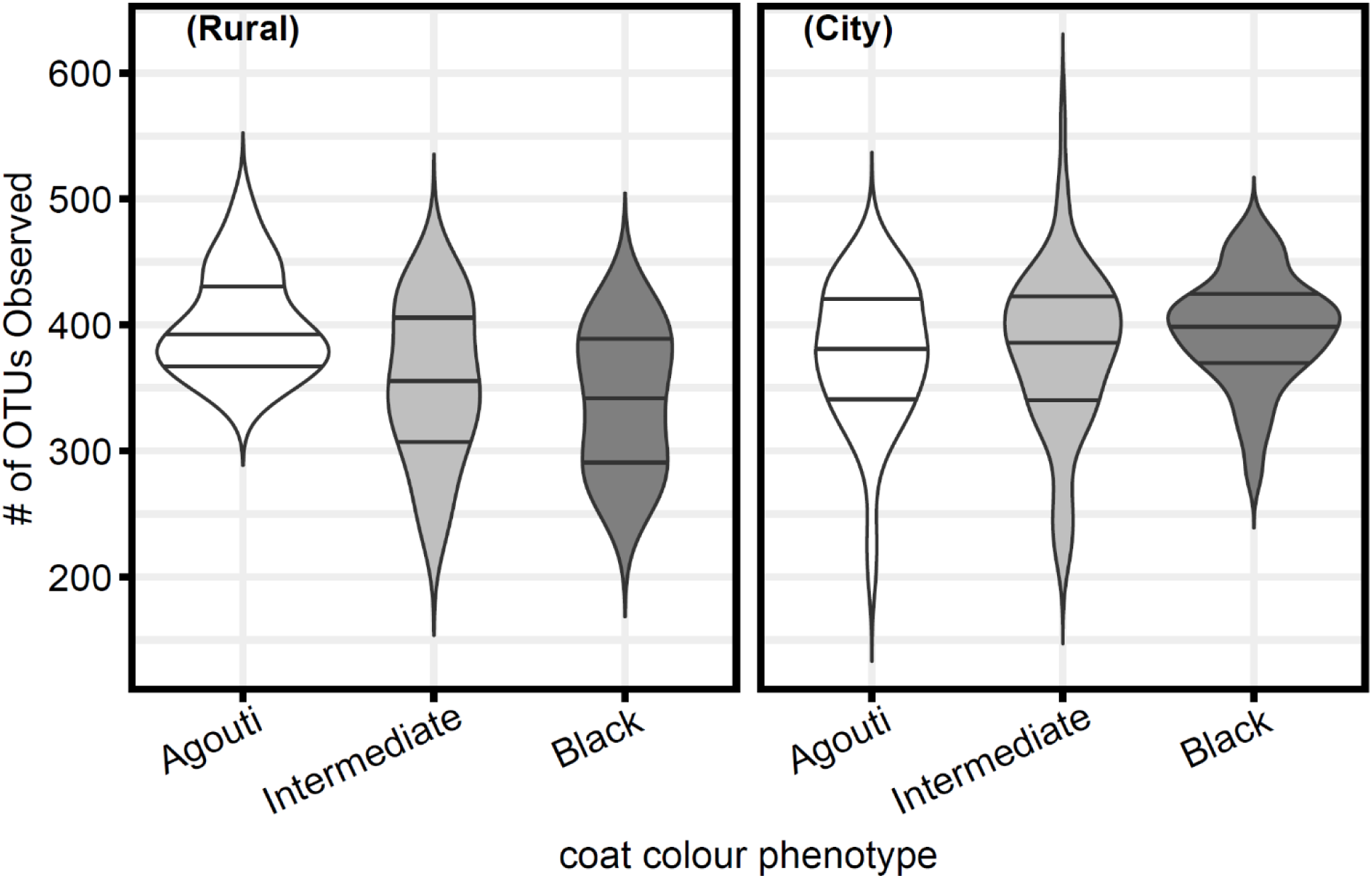
Bacterial α-diversity in the squirrel microbiome is negatively correlated with melanism at rural sites, but positively correlated with melanism among squirrels within city sites.

A similar pattern was observed with respect to the most (relatively) abundant family, *Lachnospiraceae* (37% ± 1% *SE* of all reads), which has been previously linked to city associated patterns in the microbiome of grey squirrels [29]. We observed no effect of sex (*β* = 0.03 ± 2 *SE*, *t* = 1.57, *p* = 0.12), however, *Lachnospiraceae* relative abundance was 18% ± 8% *SE* lower among city agouti squirrels than rural agouti squirrels (*t* = −2.23, *p* = 0.03). Again, we also observed a phenotype x environment interaction, whereby *Lachnospiraceae* abundance increased by 8% ± 3% *SE* for every increase in melanism mutation number at city (but not rural) sites (*t* = 2.34, *p* = 0.02; Figure 3). We note that this result should be interpreted with caution, since the inherent compositionality of amplicon datasets means that this pattern might derive from taxa other than *Lachnospiraceae*. We probe taxa specific patterns more deeply below, but regardless of the proximate response, this result highlights the effect of environment x phenotype interactions on the squirrel microbiome.

**Figure 3:**
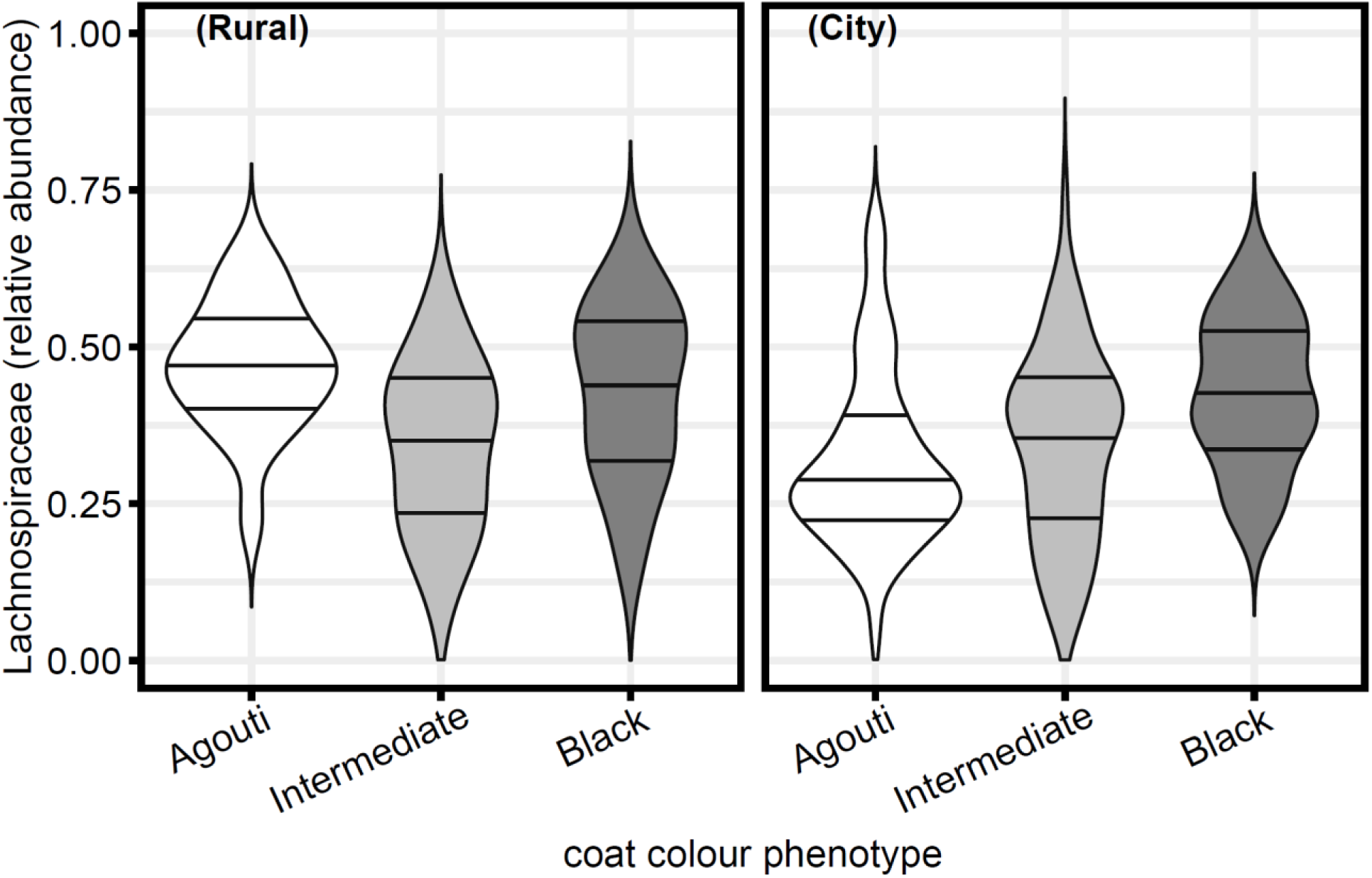
*Lachnospiraceae* relative abundance was greater among squirrels in the rural environment than those within cities, but *Lachnospiraceae* relative abundance was positively correlated with squirrel melanism among city squirrels.

### β-Diversity

OTU composition (Euclidean distance of centre log-transformed OTU counts) of the microbiome was correlated with environment (city versus rural forest; *F* = 6.96, *R^2^* = 0.03, *p* < 0.01), within-city landclass (urban campus, suburban park, urban forest, rural forest; *F* = 4.95, *R^2^* = 0.05, *p* < 0.01), trapping site (*F* = 2.78, *R^2^* = 0.08, *p* < 0.01), and sex (*F* = 1.78, *R^2^* = 0.01, *p* < 0.01) based on PERMANOVA testing. Conversely, we observed no effect of coat colour phenotype (*F* = 1.01, *R^2^* = 0.01, *p* = 0.38) and no interaction between environment and coat colour (*F* = 1.07, *R^2^* = 0.01, *p* = 0.21). Based on ordination of the bacterial microbiome (Fig 4A) and hierarchical clustering analysis (Fig 4B), squirrels from suburban parks within the city of Guelph clustered most closely with those from university campuses (built-environment sites). Conversely, squirrels from urban forests in Guelph clustered most closely with squirrels from distant rural forests (forest sites).

**Figure 4:**
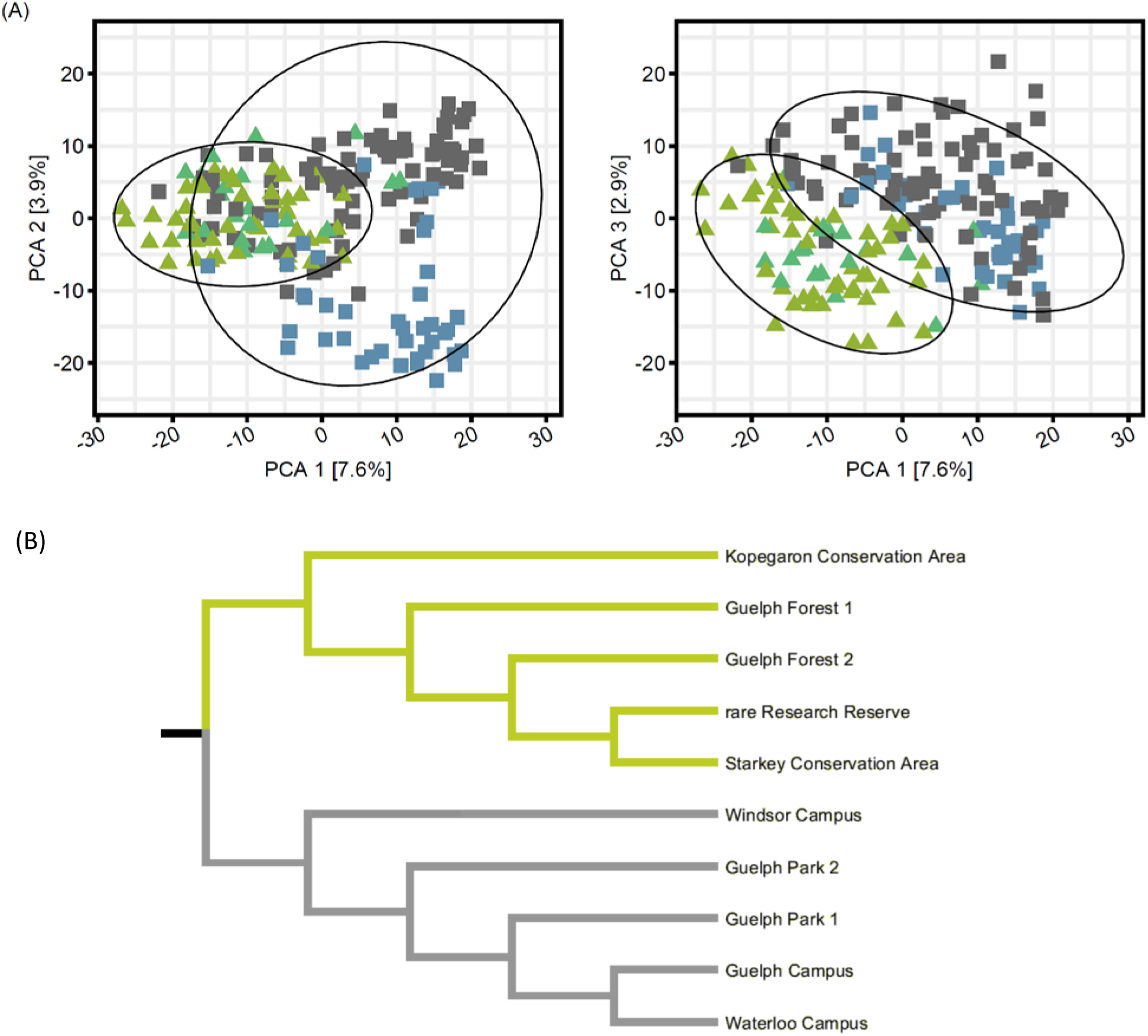
Euclidean distance (centred log-ratio transformed dataset) visualized as: (A) an ordination of the 1^st^, 2^nd^, and 3^rd^ axes of a PCoA coloured by landclass (campus: ●; suburban park: ●; urban forest: ●; rural forest: ●) and shaped by habitat type (built-environment: ▪; forest: ▴) with 95% confidence ellipses and (B) a hierarchical clustering diagram of site pooled samples of the eastern grey squirrel bacterial microbiome, green branches denote forest sites while grey branches denote built-environment sites.

Like OTU composition, phylogenetic structure of the squirrel microbiome (weighted UniFrac distance) was correlated with environment (*F* = 4.24, *R^2^* = 0.02, *p* < 0.01), within-city landclass (*F* = 5.05, *R^2^* = 0.05, *p* < 0.01), trapping site (*F* = 3.26, *R^2^* = 0.09, *p* < 0.01), and sex (*F* = 2.47, *R^2^* = 0.01, *p* < 0.01). Coat colour phenotype was marginally non-significant (*F* = 1.33, *R^2^* = 0.01, *p* = 0.06), and interestingly, we observed a significant interaction between environment type and coat colour phenotype (*F* = 1.49, *R^2^* = 0.01, *p* = 0.02).

Visualization of weighted UniFrac ordination plots suggest that the environment x phenotype interaction we observed might be partly the result of differences in β-dispersion between phenotypes across environments, rather than solely mean community dissimilarities (Fig 5A). This interpretation was supported by post-hoc permutation tests of multivariate homogeneity for phenotype-environment grouping beta-dispersion. Specifically, city agouti squirrels harboured microbiomes which were more phylogenetically variable than rural agouti squirrels (p = 0.05). No other phenotype-environment groups differed in beta-dispersion, but city intermediate squirrels trended towards displaying greater phylogenetic variability than rural agouti squirrels (p = 0.07).

**Figure 5:**
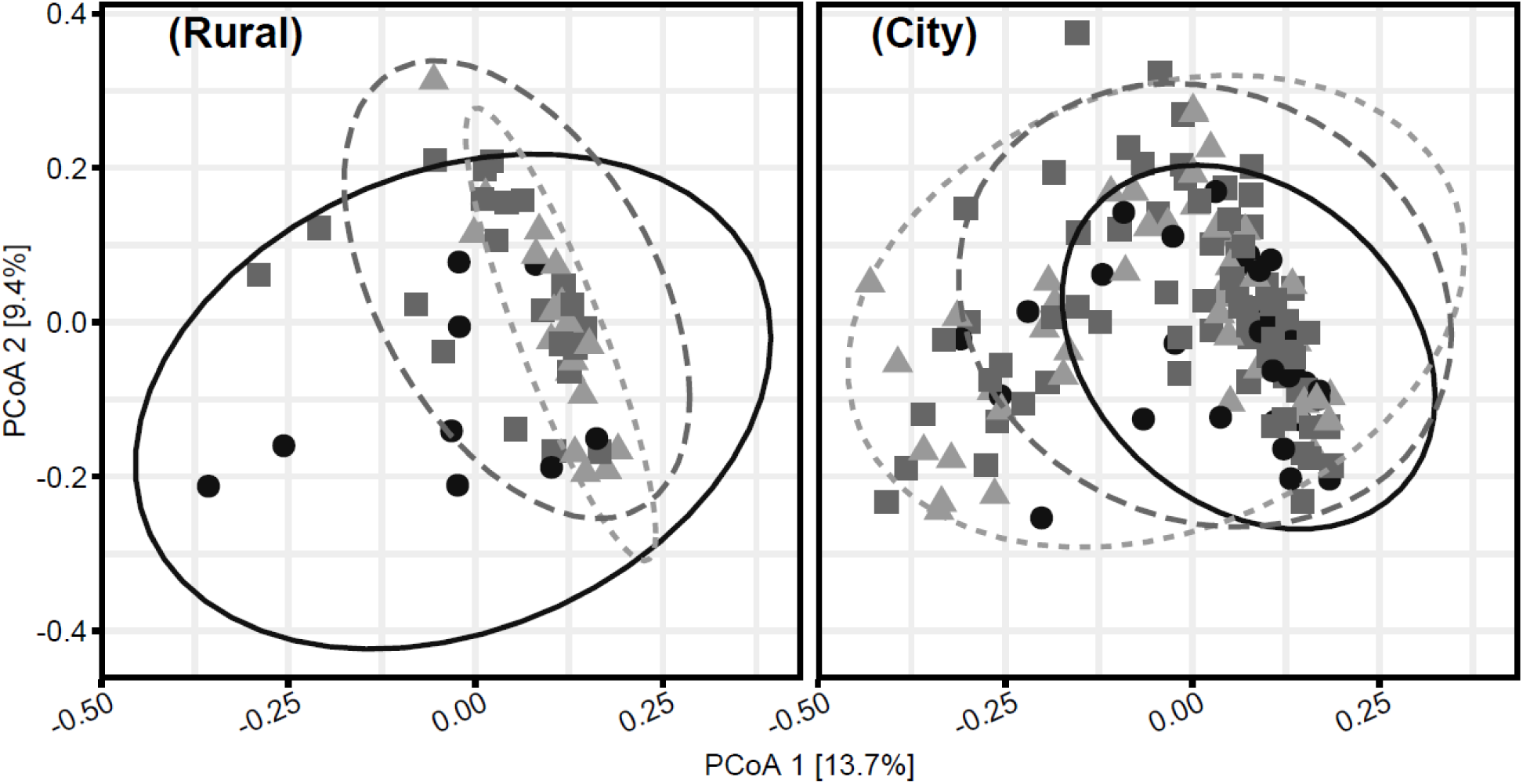
Weighted UniFrac PCoA ordination of the eastern grey squirrel microbiome separated by environment and coloured and shaped by phenotype (agouti: **▴**; intermediate: ▪; black: **●**) with 95% confidence ellipses (agouti: **---**; intermediate: **− −;** black: —). Agouti squirrels display greater beta-dispersion within cities.

### Null-Modelling Inference

Conventional β-diversity analyses are used to test whether community structure differs between groups but provide little evidence with which to infer why communities may differ. Further, patterns in β-diversity can be influenced by imbalances in α-diversity between communities [48]. Phylogeny-independent and phylogeny-weighted null modelling methods can be used to determine whether community composition (or between community dissimilarities) deviate from random expectations, given community α-diversity and pre-defined pool of γ-diversity (see Methods)[45, 47].

We tested for a phylogenetic signal among the bacteria observed in the squirrel microbiome with respect to their average relative abundance within city and rural squirrel populations. We observed a significant positive signal over short phylogenetic distances (Figure S1). This suggests that closely related bacterial OTUs tended to occupy a similar ecological niche, a result which supports the use of mean nearest taxon distance (MNTD), a measure of phylogenetic clustering/dispersion within communities [46].

Effect size standardized and box-cox transformed (λ = 0.4) absolute MNTD_ses_ values were closer to stochastic expectations (i.e., 0) among city agouti squirrels compared to rural agouti squirrels (*β* = −1.28 ± 0.58 *SE*, *t* = −2.22, *p* = 0.03); however, we observed an interaction whereby—among city squirrels—|MNTD_ses_| increased with melanism mutation number (*β* = 0.51 ± 0.24 *SE*, *t* = 2.15, *p* = 0.03; Figure 6A). Conversely, we observed no effect of sex (*β* = 0.17 ± 0.14 *SE*, t = 1.20, p = 0.23) or melanism among rural squirrels (*β* = −0.29 ± 0.21 *SE*, t = −1.38, p = 0.17).

**Figure 6:**
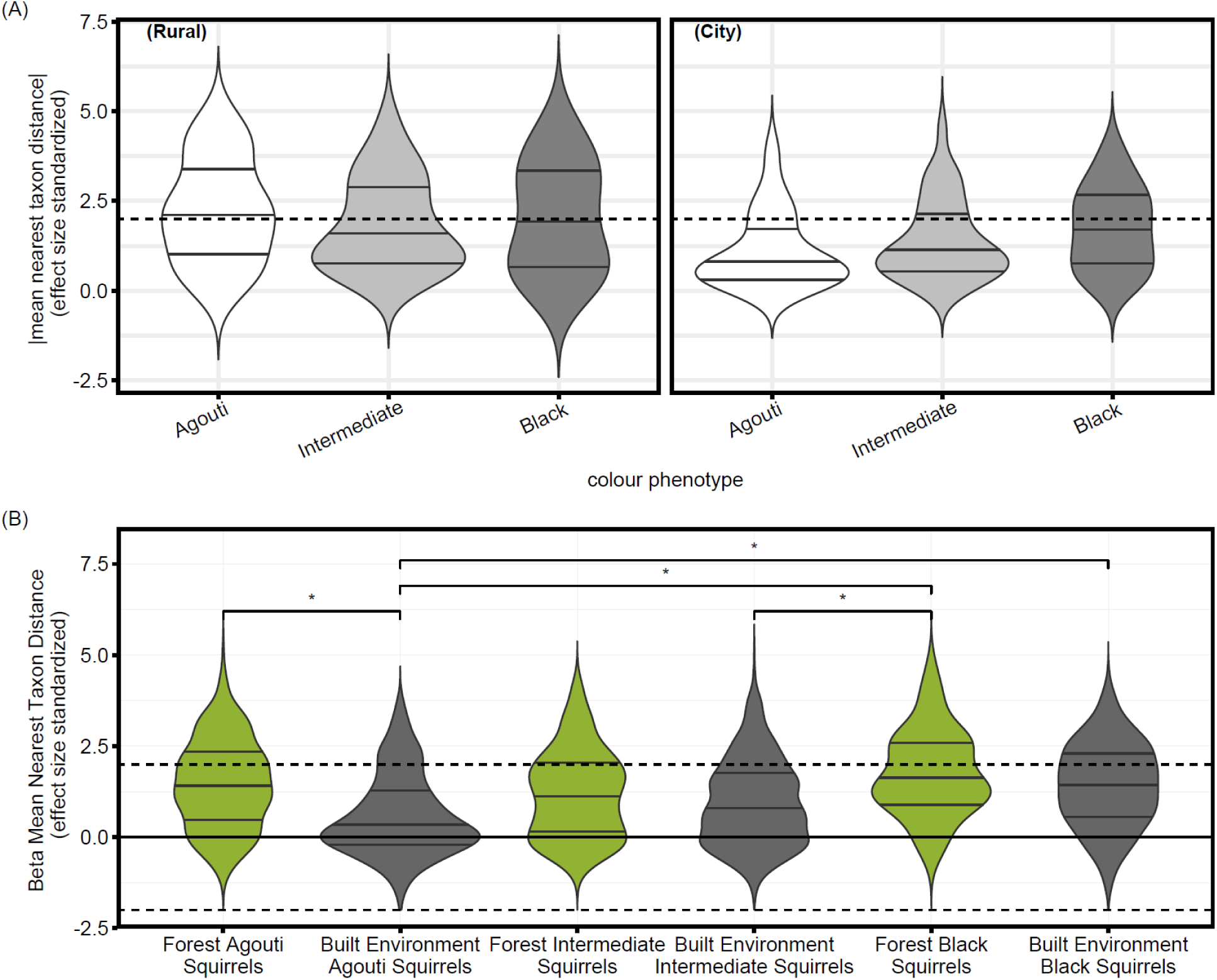
An interaction between coat colour phenotype and environment affected (A) |mean nearest taxon distances| (box-cox transformed) (city versus rural), and (B) β mean nearest taxon distance (βMNTD_ses_) (built-environment versus forest), such that city and built-environment agouti squirrels had values closer to stochastic expectations. Dotted horizontal lines denote values beyond two standard deviations from the null distribution. Horizontal brackets and ‘*’ denote significant differences in βMNTD_ses_ variance homogeneity using permutation tests.

The same principles which underlie null modeling of within community phylogenetic structure can be applied to between community contrasts. Using this approach, phylogenetic distances between all taxa in one community are measured to their nearest related taxon in another community [45]. When effect-size is standardized by the standard deviation of the corresponding null distribution, these values provide an indication of whether any two communities are more (βMNTD_ses_ > 2), less (βMNTD_ses_ < 2), or no more (|βMNTD_ses_| < 0) phylogenetically disparate than expected by chance [45]. If taxon niche-space is shallowly phylogenetically conserved (an assumption supported by the positive phylogenetic signal described above), deviations from null expectations are one indication that similar (or disparate) selective pressures act between communities. Based on a PERMANOVA test, microbiome βMNTD_ses_ values were not correlated with environment (*F* = 1.65, *R^2^* = 0.01, *p* = 0.10), within-city landclass (*F* = 0.42, *R^2^* < 0.01, *p* = 0.96), trapping site (*F* = 1.27, *R^2^* = 0.04, *p =* 0.16), sex (*F* = 1.16, *R^2^* = 0.01, *p* = 0.33), or coat colour phenotype (*F* = 1.38, *R^2^* = 0.01, *p* = 0.17); however, we observed an interaction between environment and coat colour phenotype (*F* = 1.57, *R^2^* = 0.02, *p* = 0.05). Post-hoc permutation tests of multivariate homogeneity indicated that city agouti squirrels had smaller βMNTD_ses_ variances than either rural agouti squirrels (p = 0.04) or city black squirrels (p = 0.02). Similarly, city squirrels of the intermediate morph had smaller variances than city black squirrels (p = 0.02). Notably, the magnitude of βMNTD_ses_ values among city agouti squirrels and city intermediate morph squirrels, were more consistent with stochastic expectations than they were homogenizing selection. Conversely, black squirrels maintained comparably large βMNTD_ses_ values, regardless of their environment.

Because squirrels from rural and urban forests clustered together in Euclidean-based β-diversity analyses, we additionally considered a PERMANOVA in which squirrel faecal samples from city forests were grouped with those from rural forests to test for an effect of urbanization at a finer spatial scale (forest versus built-environment). Again, we observed no effect of within city land-class (*F* = 0.00, *R^2^* = 0, *p* = 0.99), trapping site (*F* = 1.27, *R^2^* = 0.04, *p* = 0.10), sex (*F* = 1.17, *R^2^* = 0.01, *p* = 0.36), or coat melanism (*F* = 1.39, *R^2^* = 0.01, *p* = 0.14); however, we did observe a significant effect of urbanization at this finer spatial scale (human built versus forest; *F* = 2.73, *R^2^* = 0.01, *p* < 0.01) and an interaction between habitat-type and coat colour phenotype (*F* = 1.86, *R^2^* = 0.02, *p* = 0.01). As above, post-hoc permutation tests of multivariate homogeneity indicated that built-environment agouti squirrels had smaller βMNTD_ses_ variances than forest agouti squirrels (*p* < 0.01), forest black squirrels (*p* < 0.01), or built-environment black squirrels (*p* < 0.01; Figure 6B) and trended towards smaller variances than forest intermediate morph squirrels (*p* = 0.06). Built-environment intermediate squirrels likewise had smaller βMNTD_ses_ values than forest black squirrels (*p* = 0.04), and although marginally non-significant, trended towards having smaller values than either forest agouti squirrels (*p* = 0.07) and built-environment black squirrels (*p* = 0.09).

Raup-Crick bray (RC_bray_) values—which in contrast to βMNTD_ses_ measurements— indicate the magnitude of OTU turnover, independent of taxa phylogenetic relatedness. In the absence of βMNTD_ses_ deviations from stochastic expectations (i.e., |βMNTD_ses_| < 2), RC_bray_ values which are higher (> 0.95) or lower (< −0.95) are suggestive of dispersal limitation and homogenizing dispersal, respectively. Conversely, values between these extremes are considered indicative of ecological drift. Of 18943 pairwise comparisons between squirrels, 14143 (75%) had |βMNTD_ses_| < 2, and of these pairs, 13851 (98%) had corresponding values of RC_bray_ > 0.95. Environment (city versus rural; *F* = 1.21, *R^2^* = 0.01, *p <* 0.01), within-city landclass (*F* = 1.78, *R^2^* = 0.02, *p* < 0.01), trapping site (*F* = 1.34, *R^2^* = 0.04, *p* < 0.01), and sex (*F* = 1.28, *R^2^* = 0.01, *p* < 0.01) were all correlated with RC_bray_ values in a PERMANOVA test. But we observed no effect of phenotype (*F* = 1.05, *R^2^* = 0.01, *p* = 0.20), and only a marginally significant interaction between environment and phenotype (*F* = 1.10, *R^2^* = 0.01, *p* = 0.06). However, this latter result might be expected, given the significant interaction of environment and phenotype on βMNTD_ses_ values.

Post-hoc Kruskal-Wallis tests indicated that RC_bray_ values were higher among comparisons made within trapping sites than between trapping sites (χ^2^ = 530.97, *p* < 0.01). Interestingly, RC_bray_ values within city sites were smaller than those observed within rural sites (χ2 = 19.77, *p* < 0.01; Figure S2A). RC_bray_ values also differed between city and rural environments with respect to between site comparisons (χ2 = 19.99, *p* < 0.01). Specifically, RC_bray_ values were smaller between city sites than between rural sites (*p* < 0.01) or between city and rural sites (*p* < 0.01). RC_bray_ values also differed by sex both within sites (χ2 = 7.83, p = 0.02; Figure S2B) and between sites (χ2 = 15.41, *p* < 0.01). Within sites, post-hoc dunn tests indicated that female-female pairs had lower RC_bray_ values than male-male pairs (*p* = 0.02) or female-male pairs (*p* < 0.01). In comparisons made between sites, female-male pairs had larger RC_bray_ values than either female-female (*p* = 0.01) or male-male pairs (*p* < 0.01).

### Differential Abundance Testing

To generate a robust understanding of which taxa drove the apparent divide in the microbiome between squirrels in forest versus built-environments, we used two different, but complimentary, statistical approaches. The first, selection balance analysis, accounts for the compositional nature of most 16S amplicon datasets by seeking the most parsimonious log-ratio of taxa that delineate two groups [49]. Results of selection balance analyses are therefore derived from parsimony and discriminative sensitivity, rather than test statistics typical of frequentist approaches. The second approach, analysis of compositions of microbiomes with bias correction (ANCOM-BC), normalizes sequence counts by a process similar to centred-log ratio transformations and applies corrections to control for false discovery rates [50].

Selection balance cross-validation identified that the apparent divide in the microbiome between squirrels in the human built-environment (university campuses and suburban parks) versus forests (urban forest and rural forests) could be most parsimoniously discriminated by a log-ratio balance of 12 genera. Squirrels from the built environment had a greater abundance of *Odoribacter*, *Oscillospira*, *Sulfurimonas*, *Ruminiclostridium* (sub-group 5), *Coprococcus* (sub-group 3), and an unclassified genus in the class *Mollicutes* RF39, relative to *Alistipes*, *Ruminiclostridium* (sub-group 1), *Rikenellaceae* RC9, *Lachnospiraceae* UCG-010, *Clostridiales* vadinBB60 group, and an unclassified genus of *Lachnospiraceae*, when compared to forest squirrels (Fig. S3A). This balance was able to discriminate between built-environment and forest samples with high sensitivity (AUC-ROC = 0.975). Similarly, at the family level, a log-ratio of *Mollicutes* RF39 : *Rikenellaceae* most parsimoniously delineated built-environment and forest squirrels (AUC-ROC = 0.808; Fig. S3B).

The results of ANCOM-BC analyses were qualitatively like those of the selection balance analysis, although 17 genera were observed to differ between forest and human built habitats. Squirrels from the built-environment had greater relative abundances of *Mollicutes* (group RF39), *Ruminiclostridium* (sub-group 5), *Prevotella*, *Parasutterella*, *Oxalobacter*, an unclassified genus of *Eggerthellaceae*, as well genera of *Ruminococcaceae* (sub-group UCG-014, sub-group UCG-008) and *Lachnospiraceae* (*Lachnoclostridium*, *Blautia*, *Eisenbergiella*, *Coprococcus* sub-group 3). Conversely, forest squirrels had greater abundances of *Alistipes*, *Clostridiales* (sub-group vadinBB60), *Lachnospiraceae* (sub-group FCS020), *Lachnospiraceae* (sub-group NK4A136), and a third unclassified *Lachnospiraceae* genus. Notably, these environmental differences appeared driven primarily by the agouti morph. After repeating ANCOM-BC analyses upon datasets subset by colour phenotype, we observed that between forest and human built-environments, 14 genera differed among agouti squirrels, 4 genera differed among the intermediate morph, and 0 genera differed among black coated squirrels (Fig. S4).

## Discussion

Urbanization clearly affected the eastern grey squirrel microbiome. This result is consistent with findings from birds [21,23–26], reptiles [30], humans [51, 52], insects [53], plants [54], and wild mammals [29]. Unlike previous studies, we demonstrate that convergence occurs across cities, but also that substantive variation exists both between and within cities. For example, the variation explained by city-scale urbanization was comparable to the variation explained by land-class heterogeneity within a single city (campuses, suburban parks, urban forests). This is consistent with other reports that fine-scale environmental variation can play a substantive role in shaping the microbiome [55, 56] and suggests the need for caution when trying to quantify an individual’s environment with simplified univariate terms, like percent impermeable landcover in the context of urban landscapes. For example, our urban forest and suburban park sites were characterised by similarly sized urban greenspaces, but squirrels from urban forests tended to cluster with squirrels from rural forests in ordination and hierarchical clustering plots, while those from suburban parks were more alike those from university campuses. Broad-scale urbanization (city versus rural) had no effect on βMNTD_ses_, however, βMNTD_ses_ values were affected by whether squirrels were from local habitats characterized as forest or human built. These phylogenetic patterns in the microbiome suggest that environmental selective pressures act on the microbiome (perhaps indirectly through host physiology) at fine spatial scales. Therefore, although we report an effect of urbanization on the microbiome, habitat heterogeneity within cities makes a representative ‘urban microbiome’ for host species unlikely.

Anthropogenic food subsidies in cities are hypothesized to underlie microbiome variation among urban mammals [24, 31], as well as reports of obesity and hyperglycaemia [57–59]. Western diet-induced obesity in mice and pigs maintained on diets low in indigestible fibre and starch (as might be expected of a western diet) are strongly characterized by blooms of *Mollicutes* [60–62]. Although some well-known *Mollicutes* are intracellular parasites, genome reconstruction of the *Mollicutes* species associated with western-diet induced obesity in mice, suggests a specialization on simple sugars [60]. Tellingly, we observed *Mollicutes* to be one of the strongest discriminating features separating built-environment squirrels from forest squirrels in both selection balance and ANCOM-BC testing. ANCOM-BC tests further identified that built-environment squirrels harboured greater relative abundances of *Parasutterella*—sister genus to the very ecologically similar *Sutterella* [63]. *Sutterella* were observed to be a strong feature characterizing the microbiome of wild red squirrels supplemented with peanut butter, a food higher in sugar and fat than natural components of the squirrel’s diet [64]. Rather than dietary fats though, *Parasutteralla* and *Sutteralla* appear to specialize on the bile acids hosts produce to solubilize fatty dietary components [65].

A shift from fibrolytic taxa, towards those known to metabolize animal fats and host-derived products (especially bile acids) generally characterized the city and built-environment squirrel microbiome. For example, *Lachnospiraceae*—a family comprised of primarily fibrolytic specialists [66]—was less abundant among city squirrels. Despite this, a handful of *Lachnospiraceae* genera were found in ANCOM-BC analyses to be more abundant in squirrels from the built-environment, but, most of these genera have also been shown to metabolize animal host derived products (*Lachnoclostridium* [67–70], *Blautia* [71–73], *Eisenbergiella* [74]). Genera within the family *Eggerthellaceae* and the genus *Odoribacter* were likewise built-environment associated—in ANCOM-BC and selection balance analyses, respectively—and likewise specialize on bile acids and other host derived products [67,75–78]. In addition to metabolizing host products, many of these genera have been implicated in western diet-related metabolic and gastro-intestinal diseases in humans (*Lachnoclostridium* [67, 79], *Blautia* [80–83], *Eisenbergiella* [79], *Eggerthellaceae* [67,77,79,84,85], *Odoribacter* [78,83,86–88]). While diet may be the ultimate cause of the built-environment versus forest squirrel microbiome divide, host physiological responses to their diet may be a complimentary, if not more proximate, mechanism. This distinction is important, since a genetic basis to host physiological responses to their diet allows for evolution in the diet x microbiome relationship [29].

The need to consider proximate host physiological mechanisms was exemplified by that observation that although urbanization had an apparent affect on the squirrel microbiome, these affects were partly mediated by squirrel coat colour phenotype. Coat melanism was negatively correlated with OTU richness in rural environments but positively correlated with OTU richness in the city. Black squirrels harboured microbiomes which were more phylogenetically dissimilar than predicted by null expectations but did not differ systematically between environments. When parsed by phenotype, no genera differed in abundance between black squirrels sampled in built-environment and forests, whereas 14 genera differed among agouti squirrels between habitats. Agouti squirrels also displayed greater phylogenetic variability in the microbiome within cities than black squirrels, and null modelling results suggest that this might be due to stochastic (rather than deterministic) processes (|MNTD_ses_| and βMNTD_ses_ values closer to 0). If diet is indeed an ultimate cause for the effects of urbanization on the grey squirrel microbiome, then its affects might be mediated by physiological differences between squirrel colour morphs.

Fur and feather melanism—like that observed in grey squirrels—is often pleiotropically linked through the pro-opiomelanocortin system to myriad physiological pathways [44]. These pathways include baseline hypothalamic-pituitary-adrenal (HPA) physiology, HPA axis reactivity to stressors, and the immune system [89–93]—each of which is a dimension of host physiology known to affect wildlife microbiomes [29,37,94]. In eastern grey squirrels, the melanism-causing MC1rR mutation [95] has been connected to behavioural and physiological differences [96, 97], most notably thermogenic physiology [39, 40]. Specifically, melanic squirrels show greater plasticity in their ability to adaptively lower their basal metabolic rate when exposed to sub-zero ambient temperatures. This gene x environment interaction may be the result of MC1R linked pleiotropy, however, more recent evidence suggests that the MC1R mutation in grey squirrels might have originated from introgression with eastern fox squirrels (*Sciurus niger*) [98]. Therefore, the MC1R mutation—and the greater physiological plasticity with which it appears correlated—might reflect more substantive underlying genetic differences between colour morphs.

Greater physiological plasticity among black squirrels might paradoxically explain why this morph tended to maintain similar microbiomes—and with similar patterns of divergence from null phylogenetic expectations—regardless of their external environment. Conversely, if agouti squirrels are physiologically inflexible, they may be incapable of acclimating to novel urban conditions or environmental stressors, and thereby lose, or relax, control of their resident microbiota. The resultant homeostatic disruption might explain the greater evidence for ecological drift within the city and built-environment squirrel microbiome. A physiological basis to the colour phenotype patterns we observed seems likely given past research [39,40,42,44]. We cannot rule-out the alternative possibility that squirrel colour morphs occupy disparate dietary niches in the city, but even cursory observation of urban squirrels makes clear that both colour morphs readily access human food subsidies. In testing the hypothesis that microbiome plasticity is adaptive for hosts in novel environments [11], we caution that it is important to parse whether observed variance in the microbiome is the result of stochastic processes (perhaps signalling a loss of host homeostatic control) or deterministic processes (mediated by plasticity in host physiology); the outcome for host health and fitness may be very different depending on process responsible for driving beta-dispersion in the microbiome [2].

Despite the emphasis we have placed on selection and ecological drift, patterns in bacterial dispersal likely also strongly contribute to the inter-environmental variation that we observed [35]. For example, bacterial dispersal limitation undoubtedly occurs between sampling locations—an interpretation supported by the smaller RC_bray_ values observed within sites than between sites. This is unsurprising as spatial structure and social structure within [33,99–102] and between [34,55,103,104] mammalian populations has been demonstrated to affect the microbiome. More surprising, was our observation that RC_bray_ values tended to be smaller within city sites than within rural sites—despite no effect of environment on βMNTD_ses_ values. This suggests that bacterial dispersal limitation might be stronger between squirrels within rural habitats versus between squirrels within city environments. Squirrel populations persist at greater densities on urban landscapes [105], which could facilitate greater microbial exchange via more frequent interactions with conspecifics [99]. Conspecific interactions are further catalyzed by spatial clustering of anthropogenic food sources (bird feeders, garbage cans, picnic areas etc.) which are known to increase rates of pathogen transmission in wildlife [106]; the same process could as easily facilitate greater exchange of commensal or mutualistic bacteria. Further, bacterial dispersal is not restricted to among hosts of the same species, but rather, are partly shaped by trophic interactions [34]. Urban food webs tend to have fewer species, and more interactions per species (i.e. greater connectivity [107]), which might help to promote the exchange of microbiota between the microbiomes of co-occurring colonizing wildlife. Greater connectivity and spatial overlap between con-and hetero-specific hosts within urban environments could facilitate more frequent microbial dispersal when compared to rural sites.

Environment dependent patterns of microbiota dispersal may likewise partly underlie beta-diversity differences in the microbiome between environments. We observed that RC_bray_ values were smaller among pairwise comparisons made between city sites than comparisons made between rural sites or between city and rural sites. Substantive differences in abiotic and biotic factors between these environments ensure that urban and rural squirrels are very likely exposed to different pools of bacterial γ-diversity. For example, urbanization has been connected to predictable biodiversity loss and landscape homogenization [108], effects which might extend to the microbial communities. Furthermore, urban biological communities are strongly shaped by human socio-cultural factors [109]. Since the cities included in our study were built in a very similar socio-cultural context, the plant and animal species within these cities (and therefore the microbiota to which squirrels are exposed) might be more similar between cities than between rural forests. Furthermore, even when plant or animal species are found in both city and rural environments, the microbiota they transmit to squirrels may differ. For example, the phyllosphere microbiota of trees—with which squirrels closely associate—are themselves affected by urbanization [28]. Finally, the near-constant exchange of people and resources between cities facilitates gene flow and prevent the genetic isolation of urban wildlife populations [110, 111]. A similar mechanism might allow for greater bacterial dispersal between cities than between isolated rural forest fragments. Although speculative, it is important to consider an organism’s broader microbial milieu when studying host-microbe symbioses, rather than lay causality solely at the feet of host diet and physiology.

Lastly, we unexpectedly observed evidence that some of the sex effects among squirrels might derive partially from patterns in microbiota dispersal. Namely, RC_bray_ values were smaller between females than between sexes or between males, despite no effect of sex on βMNTD_ses_ values. These differences could derive from behavioural differences in bacterial transmission, or physiological differences which effect colonization success, as suggested by researchers who characterized a similar pattern of female-biased bacterial transmission among co-housed common marmosets (*Callithrix jacchus*; [112]). Among North American red squirrels, inter-individual bacterial dispersal appears to occur primarily through the maternal line [64]; therefore, a pattern of lower OTU turnover might have been observed between females because bacterial dispersal occurs both from a female’s parents and to a female’s offspring. By contrast, males may not directly contribute microbiota to their offspring. These familial-structured bacterial dispersal patterns are likely continually reinforced among related female grey squirrels, which show a greater propensity for social grooming and nest sharing when compared to males [113]. Interestingly, the opposite patterns are observed among semi-feral welsh ponies, in which males show greater centrality in both social and (inferred) microbiota dispersal networks [100]. Therefore, while dietary and physiological differences between hosts affect microbiota colonization success and abundances, organismal behaviour and variation in social structure shape microbiota metacommunities, and determine which microbiota are afforded the opportunity to colonize a new host [35,101,114].

### Summary

Urbanization affected the eastern grey squirrel microbiome at landscape and intra-city spatial scales. Characteristics of the urban squirrel microbiome echo those reported in humans and laboratory mice maintained on high-fat, low-fibre, westernized diets; however, the response of the squirrel microbiome to urbanization was coat colour phenotype dependent. Namely, the inter-environmental differences we observed were driven primarily by the agouti morph. Rather than the result of divergent selective pressures however, the city agouti squirrel microbiome displayed phylogenetic patterns more consistent with stochastic processes than black squirrels. Given the putative importance of a westernized diet in shaping the urban wildlife microbiome, colour polymorphic grey squirrels provide an interesting free-living system with which to study the gene x diet x microbiome interactions. Further research and a full holo-omics approach are required to probe the causes and fitness consequences of metagenomic plasticity and understand its importance for host species as they endeavour to colonize a new ecological niche.

## Methods

### Study Sites

Sampling was divided among three site pairs throughout southern Ontario. Site pairs were comprised of one city and one nearby rural deciduous forest outside of city limits. Each city and rural site within a pair were trapped consecutively to limit the effect of seasonal confounds. Cities are inherently environmentally heterogeneous, therefore, to standardize our efforts, we limited sampling to urban university campuses as our representative city sites. Those sites were the University of Guelph (43°31’52.45″N 80°13’36.70″W), University of Waterloo (43°28’17.68″N 80°32’36.03″W), and University of Windsor (42°18’23.77″N 83°4’1.92″W), which were paired with forests at Starkey Hill Conservation Area (43°32’37.99″N 80°9’15.82″W), the rare Charitable Research Reserve (43°22’57.72″N 80°20’54.43″W), and the Kopegaron Woods Conservation Area (42° 4’38.52″N 82°29’34.04″W), respectively. To understand how habitat heterogeneity within cities can affect the microbiome, we also sampled other urban land-classes within the City of Guelph: two suburban parks (Exhibition Park and Royal City Park) and two urban forests (the University of Guelph’s Dairy Bush and Arboretum)—small forest fragments entirely enveloped by an urban landscape.

### Capture Protocol

All trapping occurred in southern Ontario between May 9th and September 12th using tomahawk Model 102 traps (Tomahawk Live Trap Co., WI, USA) baited with peanuts. Traps were set between 6:00-16:00 and checked in 1hr intervals. Upon capture, we transferred squirrels to a cloth bag and checked in 15min intervals for faecal pellets. Using single-use sterilized toothpicks, we collected fresh faecal pellets which we stored on ice in the field before transfer to −20°C for storage within 4hrs of collection. After faecal pellet collection, we transferred squirrels to a handling bag, recorded sex, and marked squirrels with unique alpha-numeric ear-tags before release to allow for identification of recaptures and prevent resampling of individuals.

### Sequencing and Bioinformatics

Using QIAamp DNA Stool Mini Kits (Qiagen, Hilden Germany), we extracted bacterial DNA from 0.2 g subsamples of faecal samples, which in other rodents are indicative of bacterial communities in the large intestines [115]. Extracts underwent triplicate PCR amplification of the 16S rRNA gene v4 region (515F-806R modified primers; Walters et al. 2015) at MetaGenomBio Inc (Toronto Canada) alongside pcr negative controls. Triplicate PCR products were then pooled prior to sequencing on an Illumina MiSeq using v2 chemistry (250bp read). We then processed pair-end reads in mothur using a standardized amplicon processing pipeline [117], aligned sequences to the silva v132 reference database [118], and classified operational taxonomic units (OTUs) clustered using OptiClust [119] based on a 97% similarity threshold. To remove potential contaminants or sequencing errors, we discarded all non-bacterial OTUs and OTUs which were not represented by at least 1 read in 5% of the samples prior to analysis [104]. In total, we generated 3,826,931 merged bacterial amplicon sequences, with an average sequencing depth of ∼20,000 reads/sample, after assembly, quality control, and filtering. Samples were rarefied to a depth of 9,697 reads (rarefaction curve: Figure S5), except for ANCOM-BC analysis and analyses pertaining to Euclidean measures of β-diversity which were calculated from raw count # centred log ratio transformed OTU datasets [120]. A relaxed neighbour-joining method was used to construct a phylogenetic tree using the mothur implementation of clearcut [117, 121]. We sequenced extraction kit negative controls which contained swabs of the bags which were used to cover the tomahawk traps. However, negative controls showed no evidence of substantive kit or field contamination.

### Diversity Testing

We tested whether microbiome α-diversity (# of OTUs) differed with sex, coat melanism mutation number (agouti = 0, intermediate = 1, black = 2), environment (city versus rural), and an interaction between coat melanism and environment using linear mixed effects models in the R package ‘lmerTest’ [122], treating sampling location as a random effect. To determine whether bacterial microbiome β-diversity differed between city and rural eastern grey squirrel populations, we used permutational multivariate analyses of variance (PERMANOVA) tests of β-diversity in Euclidean and weighted UniFrac space. PERMANOVAs were parameterized by environment type (city versus rural), landclass (campus, suburban park, urban forest, rural forest), sampling location, sex, coat colour phenotype, and an interaction between phenotype and environment type, in that order. To test for differences in environment-phenotype groupings, we used permutation tests of multivariate homogeneity, with β-dispersion calculated relative to the group centroid with a bias adjustment for small sample sizes.

### Community Null Modeling

To determine whether observed patterns of microbiome diversity deviated from stochastic expectations, we used a null modelling approach [45,46,123]. These methods are predicated on species’ ecological niche-spaces being non-independent of their phylogeny [47], a notion broadly supported via analysis of functional traits derived from publicly archived microbial genome sequences [124, 125]. We tested for a positive phylogenetic signal in urban associated niche-space using the *phyloCorrelogram()* function from the R package ‘phylosignal’ [126]. OTU niche space was estimated as the deviation of an OTU from a 1:1 line of mean OTU relative abundance in city versus rural habitats (Figure S1).

In brief, we calculated mean nearest taxon distance (MNTD) and β mean nearest taxon distance (βMNTD), which are among community and between community measures of phylogenetic dissimilarity, respectively. For all analyses, these measures were effect size standardized by the mean and standard deviation of MNTD/βMNTD null distributions created through stochastically assembled communities which possessed the same α-diversity as the focal observed communities [45, 46]. Null distributions were created by randomly shuffling taxa names and relative abundances across the system’s γ-diversity phylogenetic tree (MNTD: 9999 iterations, βMNTD: 999 iterations) using the function *ses.mntd()* from the package ‘picante’ [127] and *ses.comdistnt()* from the package ‘MicEco’, respectively [128]. Additionally, we calculated RC_bray_ values, a phylogeny-independent measure of between community OTU turnover [46, 48]. In brief, observed Bray-Curtis values were compared to 9999 probabilistically assembled community pairs of the same α-diversity as observed community pairs [46].

A linear mixed effects model was used to test for an effect of sex, environment, and colour phenotype, as well as a phenotype x environment interaction on |MNTD|, which we used as a measure of among community deviation from stochastic expectations. Again, sampling location was included in the mixed model as a random effect. To determine whether the phylogenetic structure between communities were more similar (βMNTD_ses_ < 0) or more disparate similar (βMNTD_ses_ > 0) than expected by chance alone (|βMNTD_ses_| > 2), we used a PERMANOVA (parameterized as in the β-diversity analyses described above). As above, post-hoc permutation tests of multivariate homogeneity were used to parse significant relationships. Dispersion in βMNTD_ses_ values were calculated relative to the group centroid with a bias adjustment for small sample sizes[129], using the ‘vegan’ function *betadisper()* [130].

RC_bray_ values were analyzed using a PERMANOVA, as described above. To make within and between group comparisons, we used post-hoc Kruskal-Wallis and dunn tests to further parse results into within versus between trapping sites. We used an ultrametric phylogenetic tree transformed using the *chronos()* function in the R package ‘ape’ (λ = 1; [131]) for all phylogeny-weighted null modelling. Unless specified, all other analyses were completed in R (v. 3.5.1) using the R package ‘phyloseq’ [132].

### Differential Abundance Testing

To identify how the bacterial microbiome differed between squirrels in forests versus the built-environment, we performed selection balance analyses after binning OTUs to genus and family [49]. Secondarily, we performed ANCOM-BC differential abundance tests and evaluated the agreement between these opposing statistical approaches [50].

## Declarations

### Ethics approval and permits

Animal utilization protocol was approved by the University of Guelph Animal Care Committee (AUP no. 3506). Trapping authorization was granted by the Ontario Ministry of Natural Resources (WSCA no. 1087323).

### Availability of data and material

All sequence data have been archived with the NCBI SRA (under embargo) and are available from the corresponding author upon reasonable request. All data and scripts will become publicly available upon publication of the peer-reviewed article.

### Authors’ contributions

AEM and MRS designed the study. MRS collected samples, completed labwork, and performed analyses. AEM and MRS wrote the manuscript.

## Acknowledgements

We thank the rare Charitable Research Reserve, the Essex Region Conservation Authority, the Grand River Conservation Authority, the City of Guelph, the University of Guelph, the University of Waterloo, and the University of Windsor for logistical support and allowing access to their respective properties. Thanks also to Michelle Hotchkiss for her aid in data collection.

---

**Figure S1:**
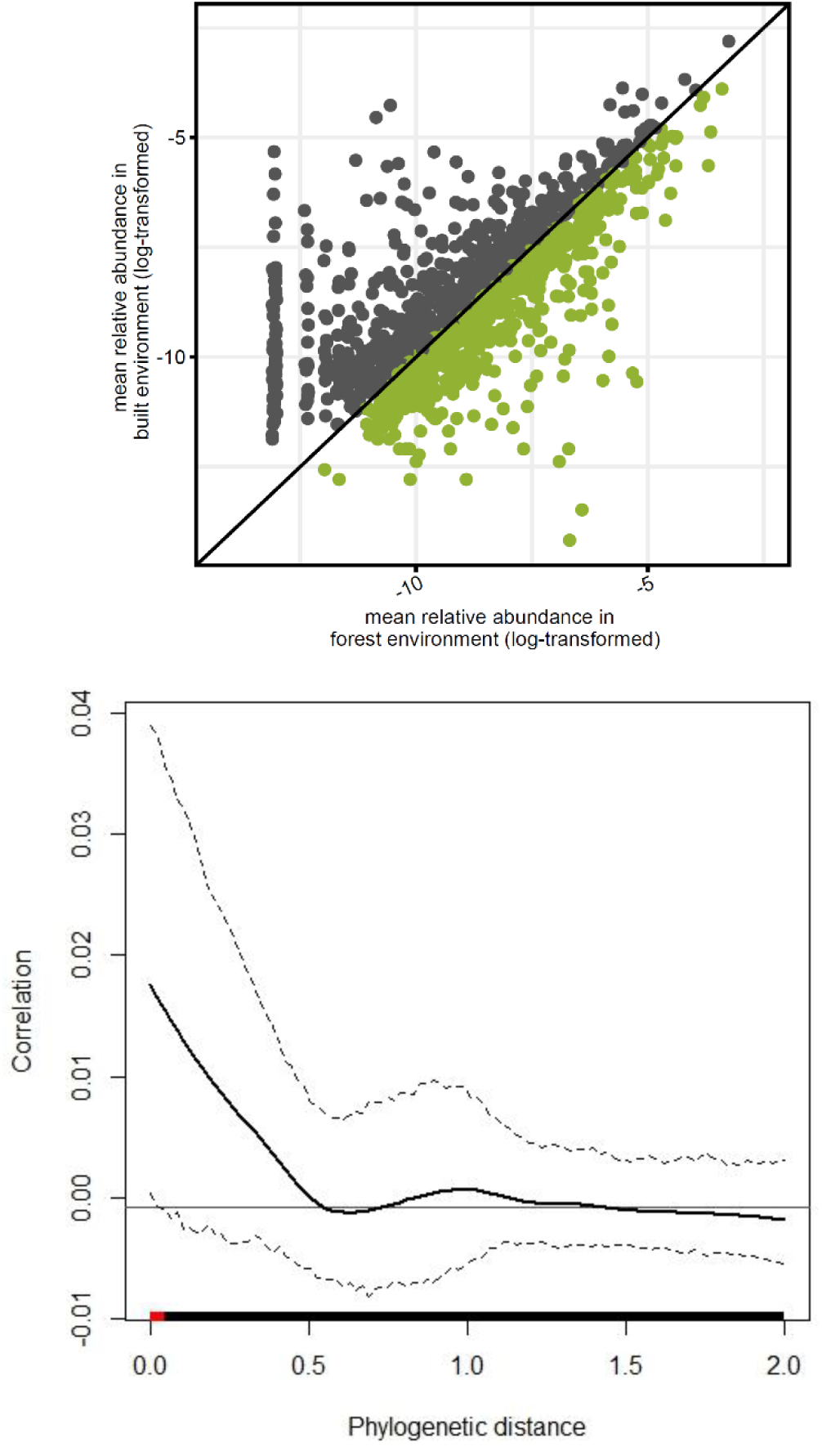
Plots of (A) log-transformed mean relative abundance of OTUs (points) among forest (●) versus built-environment (●) squirrels with 1:1 line (solid black) and (B) a phylocorrelogram of OTU deviation from the 1:1 line, which shows a positive phylogenetic signal over short phylogenetic distances (significance identified by a red line) for bacterial OTU bias towards squirrel local environment type.

**Figure S2:**
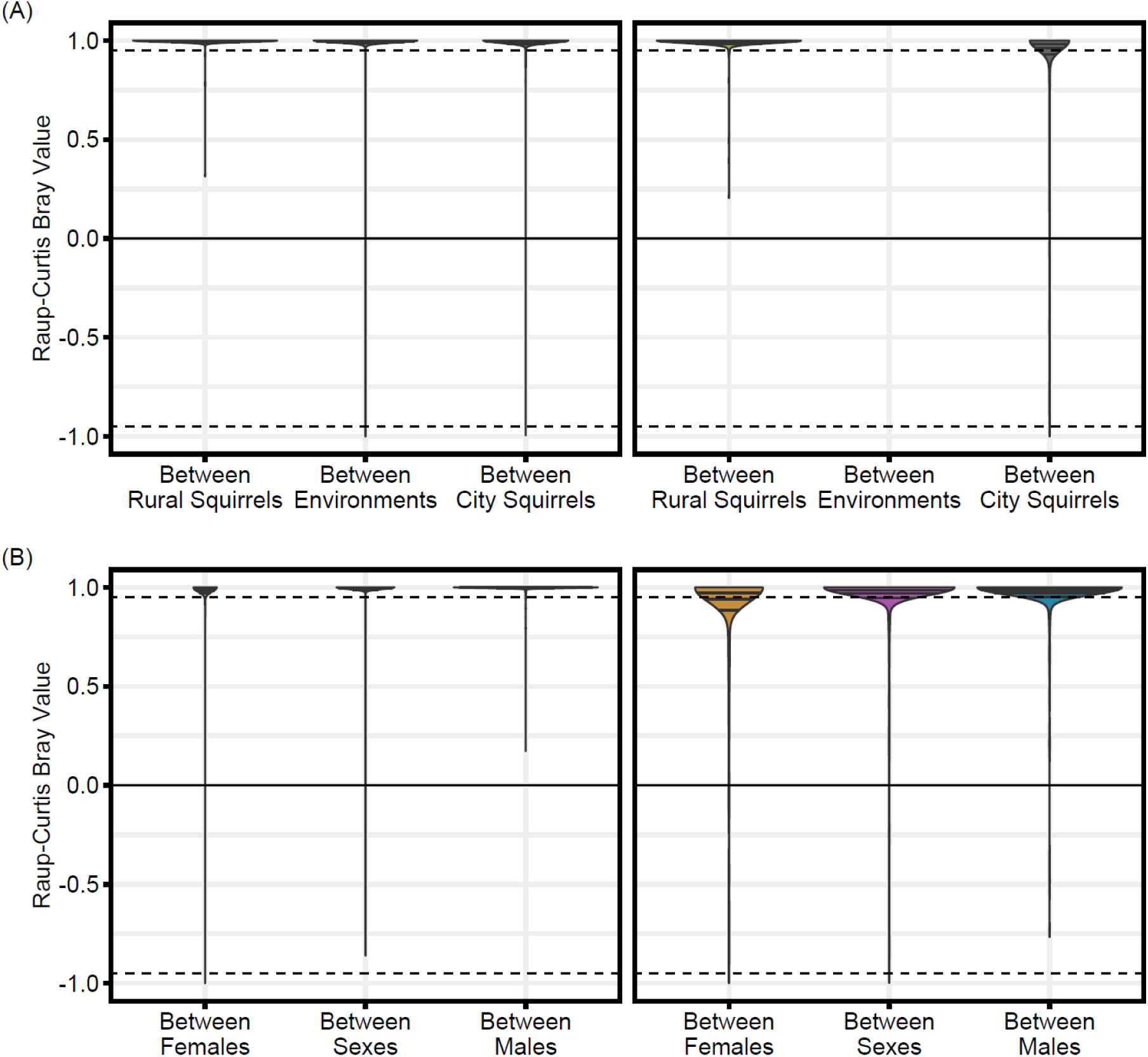
Plots of RC_bray_ values among and between (A) environments and (B) sexes, facetted by pairwise comparisons made between sites (left) verses within sites (right).

**Figure S3:**
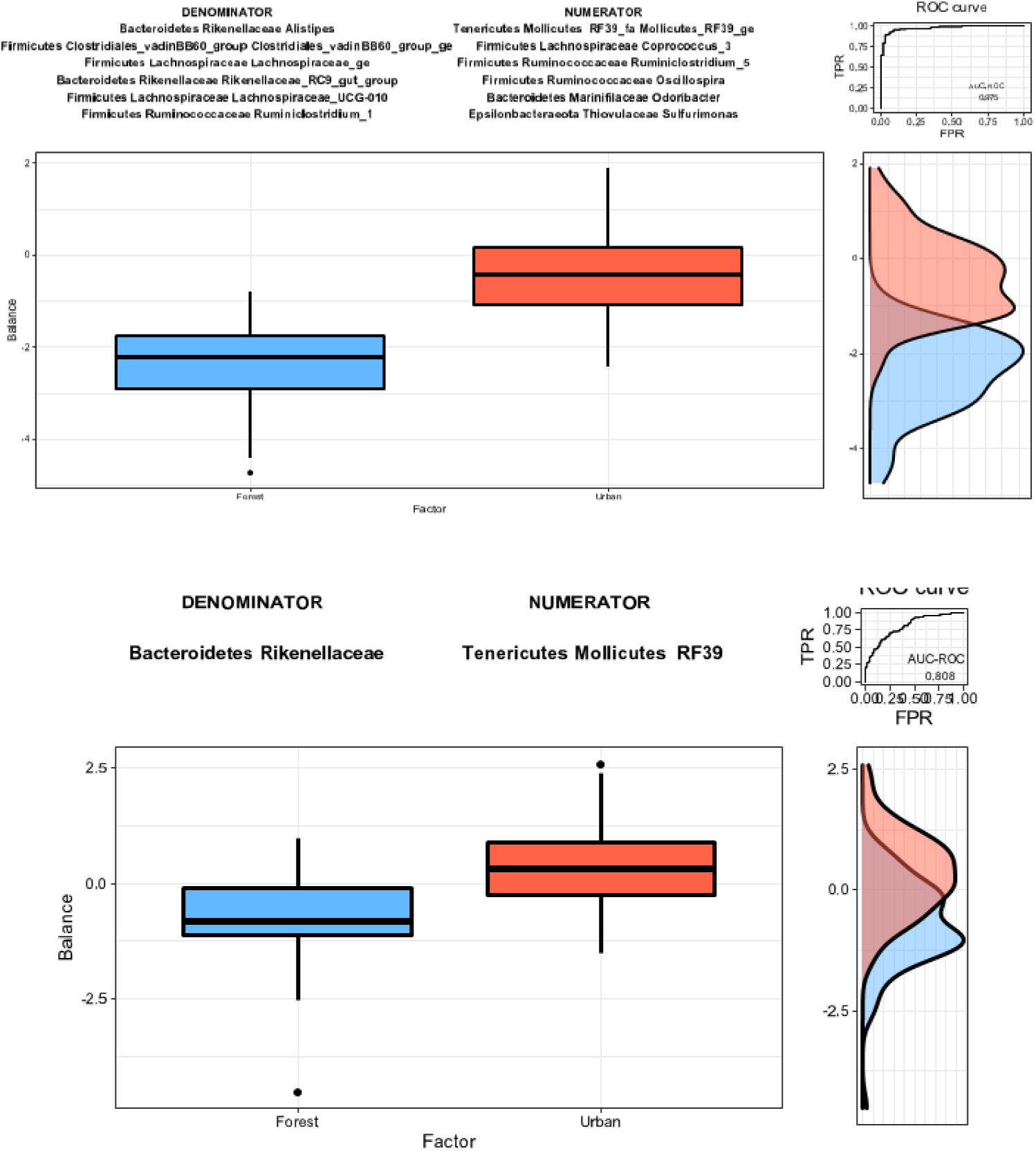
Boxplot, AUC-ROC curve, and density plots from selection balance analyses used to discriminate between forest and urban microbiomes at the bacterial taxonomic level of a) genus and b) family.

**Figure S4:**
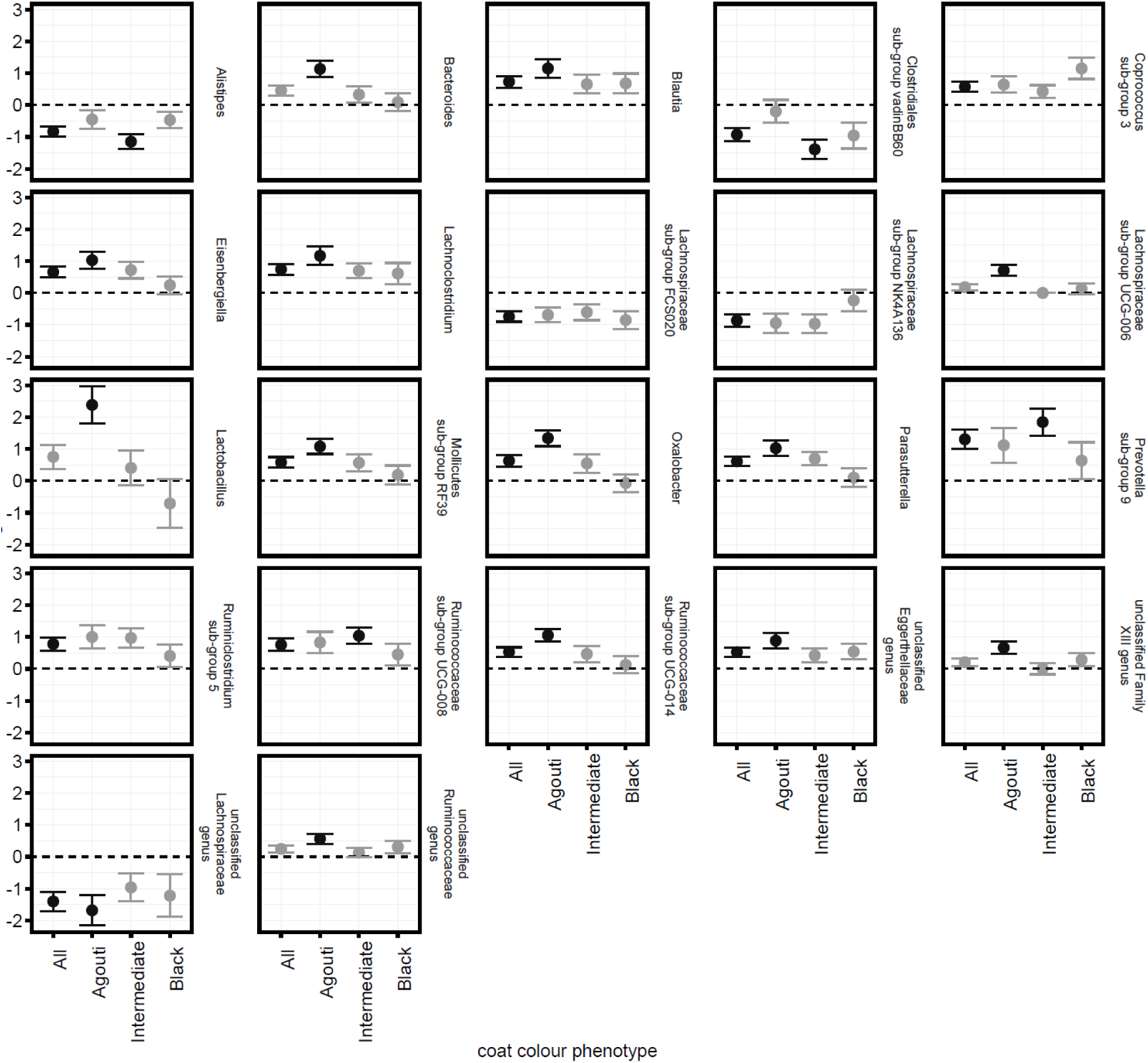
Forest plots of ANCOM-BC estimated log difference (dot) and standard error (whiskers) of genus abundances which significantly differed (black) or did not differ (grey) between forest (y-valus <0) and built-environment (y-values >0) squirrels of pooled, or parsed coat colour phenotypes.

**Figure S5:**
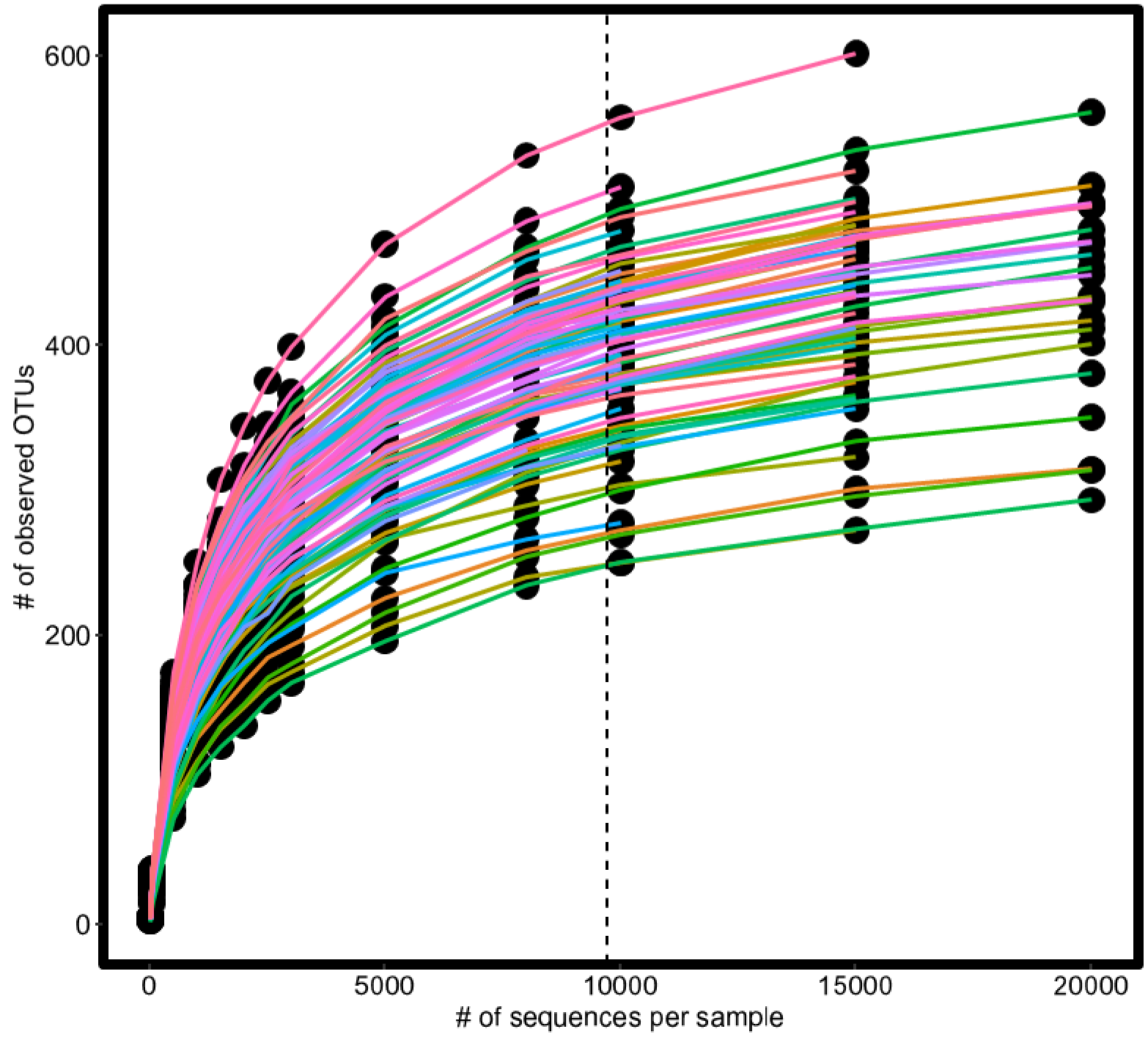
A rarefaction curve of observed # of OTUs versus sub-sampled sequencing depth. Coloured lines represent separate individuals. The dotted line demarcates level of rarefaction used for analyses.

